# Identification of a new Corin atrial natriuretic peptide-converting enzyme substrate: Agouti-signaling protein (ASIP)

**DOI:** 10.1101/2023.04.26.538495

**Authors:** Bryan T. MacDonald, Nadine H. Elowe, Colin W. Garvie, Virendar K. Kaushik, Patrick T. Ellinor

## Abstract

Corin is a transmembrane tethered enzyme best known for processing the hormone atrial natriuretic peptide (ANP) in cardiomyocytes to control electrolyte balance and blood pressure. Loss of function mutations in Corin prevent ANP processing and lead to hypertension. Curiously, Corin loss of function variants also result in lighter coat color pigmentation in multiple species. Corin pigmentation effects are dependent on a functional Agouti locus encoding the agouti-signaling protein (ASIP) based on a genetic interaction. However, the nature of this conserved role of Corin has not been defined. Here we report that ASIP is a direct proteolytic substrate of the Corin enzyme.

## Introduction

The atrial natriuretic peptide-converting enzyme Corin is a complex membrane bound protease (1). Corin is a type II transmembrane protein with an extensive extracellular region containing two Frizzled domains, eight LDLR type A repeats, and a Scavenger-like receptor domain, which may collectively contribute to substrate recognition. In the heart, release of pro-ANP is controlled by blood volume and locally processed by Corin via its carboxyl terminal Trypsin-like catalytic domain (2). Corin cleaves the pro-ANP cardiac prohormone resulting in a 28 amino acid mature ANP capable of stimulating the NPR-A natriuretic peptide receptor to activate cGMP signaling pathways and regulate vasodilation (3). Consequently, loss of function human variants in the Corin/ANP/NPR-A pathway result in hypertension (4). The Corin deficient knockout mouse model is viable and displays a similar hypertensive phenotype due to the inability to process pro-ANP into ANP (5). It was also found that Corin knockout mice have lighter hair pigmentation in the presence of a functional Agouti locus indicating an additional role for Corin regulating coat color (6).

At the surface of the melanocyte, alpha melanocyte-stimulating hormone (αMSH, encoded by pro-opiomelanocortin, Pomc) binds with high affinity to the melanocortin receptor MC1R to activate cAMP signaling, resulting in the production of eumelanin (dark pigment). This pathway can be blocked by the secreted ASIP which binds the MC1R receptor as an inverse agonist to decrease cAMP signaling and promote the production of pheomelanin (yellow pigment). Both MC1R (*e, extension*) and ASIP (*a, agouti*) are well known for their classic coat color genetics (7). Mice with ASIP loss of function mutations revert to a dark black eumelanin coat color (8). A similar dark pigmentation phenotype was found in ATRN mutants (*mg, mahogany* or attractin), shown to be a co-receptor for ASIP to facilitate MC1R inhibition (9). Conversely, overexpression of ASIP or its paralog Agouti-related protein (AgRP) results in pheomelanin production with a phenotypic light yellow coat color (10).

In the wild, mice and many mammals have a mottled brown appearance arising from a band of yellow pigment in the hair shaft (11). This is generated by a regulated pulse of ASIP expression to alternate pigmentation within an individual hair shaft (12). The Corin knockout mouse has a wider yellow band of pheomelanin pigmentation, which would be consistent with more ASIP activity during generation of the hair shaft (6). Corin missense and loss of function variants have been reported in the yellow coated Hanwoo Korean cattle (13), the golden Bengal tiger (14), the domestic Siberian golden tabby cat (15), the extreme-sunshine tabby cat (16) and the copper coated British recessive *wideband* cat (17); all with expanded lighter color pigment in their hair shafts. These various natural *wideband* Corin mutants bear a similarity with a targeted catalytic point mutation in the mouse demonstrating that the genetic interaction is tied to Corin enzymatic activity for coat color regulation (18). However, a clear connection between Corin and ASIP at the protein level has not been fully described. Here we definitively show ASIP is substrate for the Corin protease.

## Materials and Methods

Purified soluble Human Corin WT (residues 124-1042), Corin S985A catalytic mutant and NPPA-FlagHis were prepared as previously described (19,20). Human NPPA (residues 26-151) or recombinant Human ASIP/Agouti Protein, untagged (R&D 9094-AG-50, residues 23-132) were used in Corin digestion reactions in a PBS buffer supplemented with 100uM Calcium. Full-length protein digestions were performed with 10uM substrates combined with 50-200nM Corin enzyme at room temperature or 37C. Corin enzyme concentrations for each experiment: 50 nM for Fig 1B; 100 nM for Fig 1C, FigS1A and Fig 3; 200 nM for Fig2 and FigS1B. Proteins in Laemmli SDS-reducing sample buffer (Boston BioProducts, BP-111R) or non-reducing sample buffer (Boston BioProducts, BP-111NR) were resolved by sodium dodecyl sulfate-polyacrylamide gel electrophoresis (SDS-PAGE) using 4-20% Mini-Protean TGX gradient gels (4561094, Bio-RAD) and stained with Coomassie Blue. Human ASIP protein was detected by western blot using an Anti-ASIP antibody, basic region epitope 56-70 (Sigma, SAB1100893). Intact mass of ASIP and ASIP digestion products was determined by LC-MS using our approach for detection and quantitation of the NPPA prodomain and ANP (21). N-terminal peptide sequencing (Edman degradation) was performed at the Tufts University Core Facility. AnaSpec custom ASIP fluorogenic peptides with N-terminal fluorescent donor and a lysine conjugated C-terminal quencher: 8-mer Hilytefluor488-LNKKSKQI-K(QXL520)-NH2, 11-mer Hilytefluor488-ALNKKSKQIGR-K(QXL520)-NH2, 15-mer Hilytefluor488-IVALNKKSKQIGRKA-K(QXL520)-NH2. Assay conditions for the fluorogenic peptide cleavage assay used 100nM Corin and 100uM substrate in PBS buffer supplemented with 100uM Calcium at room temperature.

**Figure 1.**
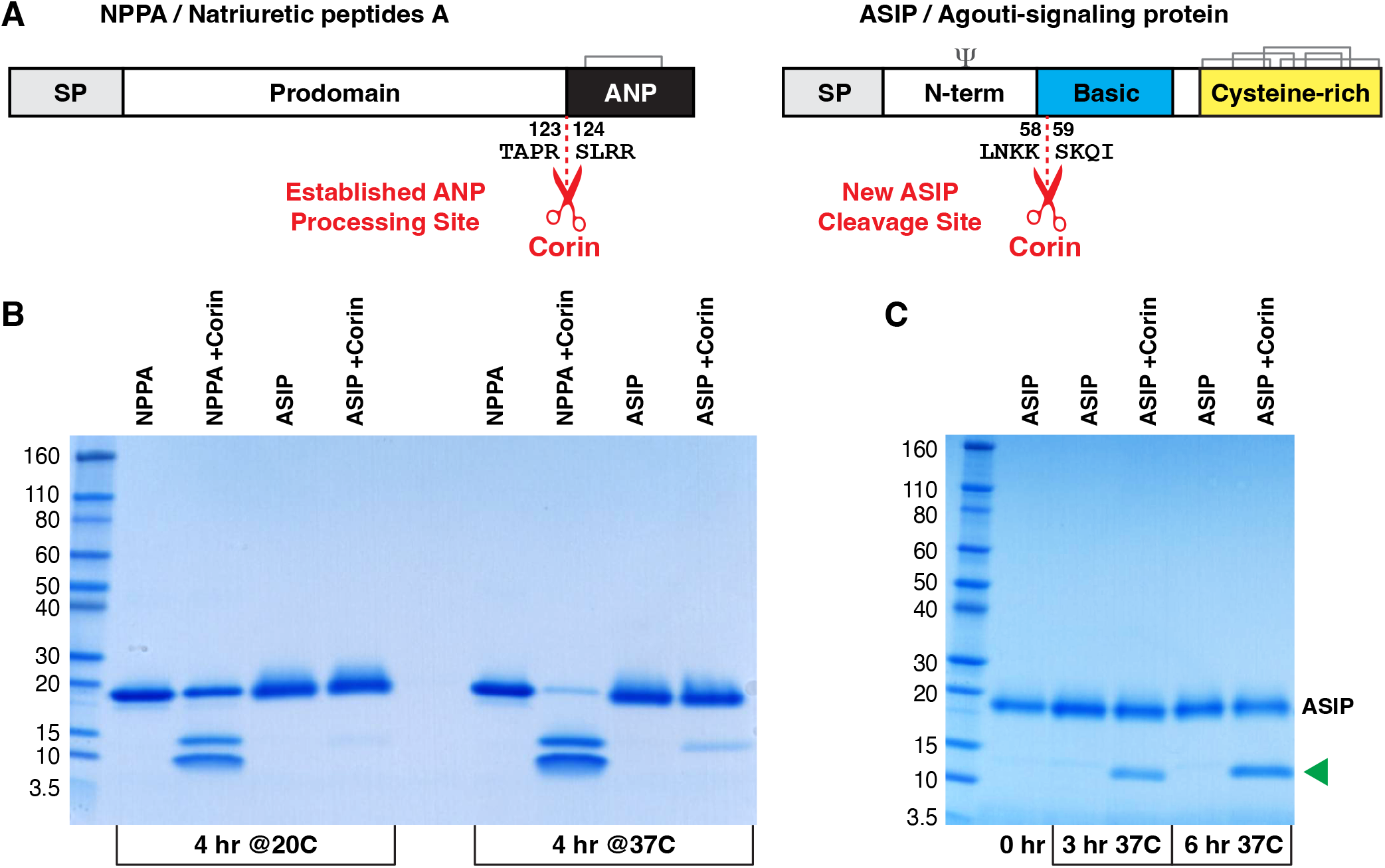
Corin dependent cleavage of NPPA and ASIP. A, Protein domains of NPPA and ASIP showing the locations of the Corin cleavage sites. SP indicates the signal peptide regions which are absent from the mature secreted proteins. Cysteine disulfide bonds and N-glycosylation are shown above the protein domains. B, Corin incubation with purified NPPA or ASIP results in activity dependent substrate cleavage, resolved by SDS-PAGE and stained with Coomassie blue. Purified NPPA contains a carboxyl terminal Flag6His tag and migrates at a similar position to the purified glycosylated ASIP. The Corin processed ANP-Flag6His product migrates faster than the NPPA prodomain. ASIP cleavage product(s) are roughly half the size of the purified ASIP under reducing conditions. C, Time dependent ASIP cleavage by Corin resolved by SDS-PAGE, new Corin dependent product(s) indicated with green arrowhead.

**Figure 2.**
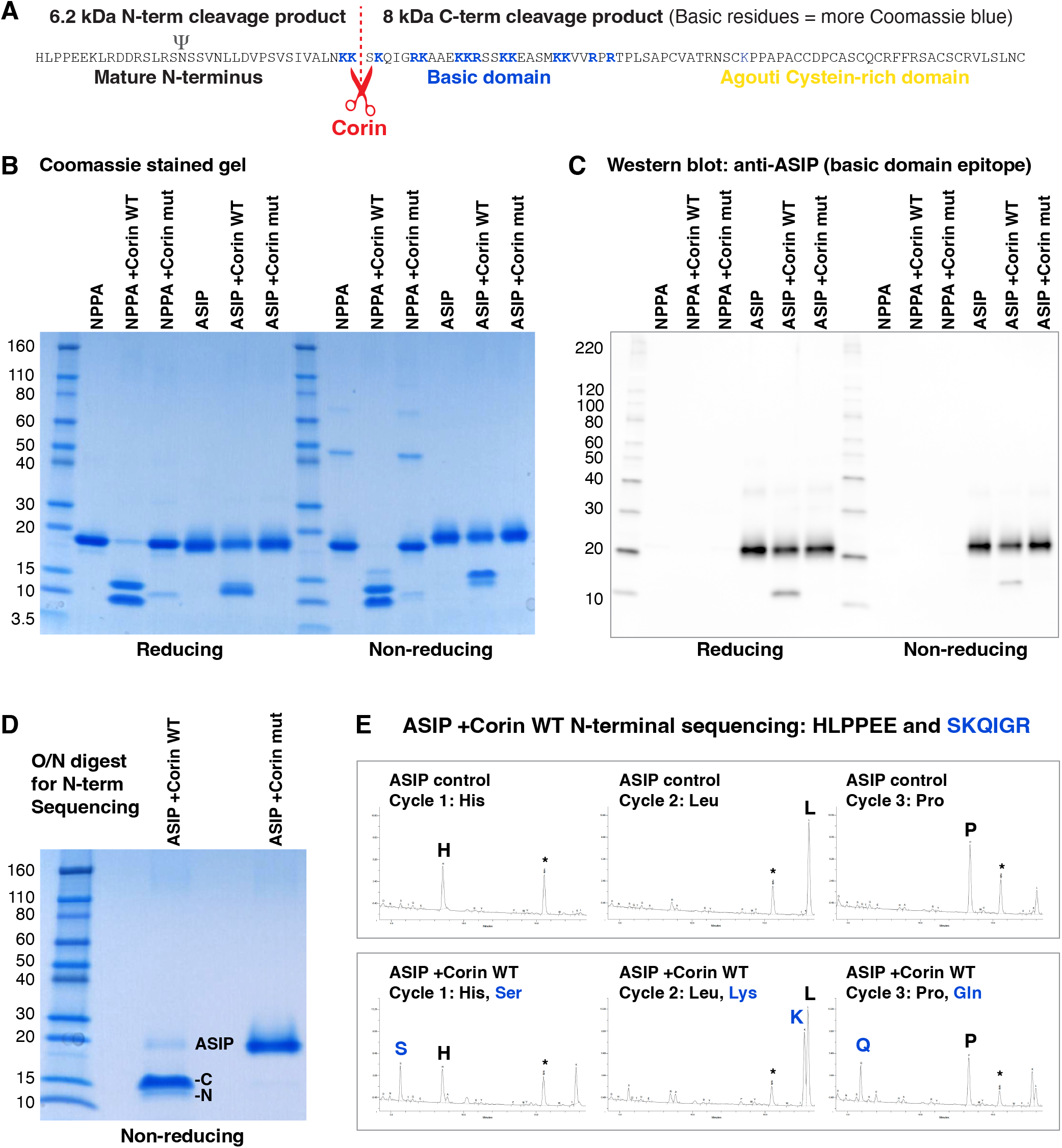
Confirmation of ASIP cleavage at the beginning of the basic domain. A, Amino acid sequence of mature human ASIP and location of the Corin cleavage site. B and C, Corin dependent cleavage reactions loaded in reducing and non-reducing sample buffers to resolve ASIP cleavage products. Corin mut contains an inactivating S985A mutation. B, Coomassie gel staining of basic residues results in a proportionally higher staining of the ASIP C-terminal cleavage product. C, Western blot using an ASIP antibody raised to a basic domain epitope detects the full-length ASIP and C-terminal cleavage product. D, Overnight Corin incubation resolved on a non-reducing gel to show ASIP cleavage, a parallel digest was submitted for N-terminal sequencing. E, N-terminal sequencing results for ASIP cleaved by Corin and HPLC-MS amino acid peaks for the first three cycles. Diphenylthiourea (dptu) peaks, a byproduct of Edman degradation, are indicated with asterisks.

**Figure 3.**
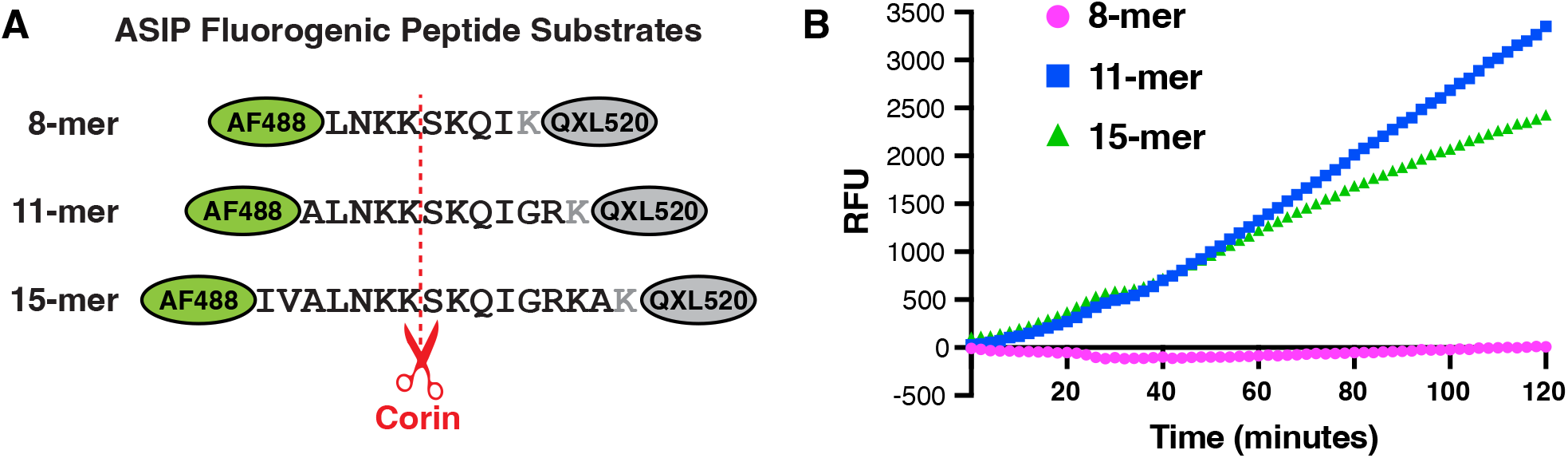
ASIP fluorogenic peptides cleaved by Corin. A, Synthetic ASIP peptides containing an N-terminal HL488 fluorescent donor and a C-terminal lysine conjugated QXL520 quencher. Corin mediated cleavage releases the quencher resulting in fluorescent signal. B, Time dependent Corin enzymatic cleavage of the ASIP 11-mer and ASIP 15-mer fluorogenic peptides, RFU relative fluorescence units.

## Results

### Purified Corin cleaves both pro-ANP and ASIP

Using a purified soluble Corin WT protein, we recently demonstrated Corin enzymatic activity in the context of a purified Natriuretic Peptide A (NPPA or pro-ANP) protein containing C-terminal Flag and His tags (20). Soluble Corin processed pro-ANP at the Arg123-Ser124 cleavage site resulting in a functionally active ANP-FlagHis (Fig1A). Incubation of soluble Corin with pro-ANP at room temperature or at 37C generated the prodomain and ANP-FlagHis products, resolved by SDS-PAGE and stained with Coomassie blue (Fig1B). Intensity of the full-length pro-ANP band diminished after incubation at 37C and the intensity of the smaller, faster migrating ANP-FlagHis band increased, demonstrating that Corin cleavage is temperature dependent. In parallel, we incubated soluble Corin with purified human ASIP protein. Following signal peptide cleavage, full-length ASIP protein contains a glycosylation site in the N-terminus, a central basic domain enriched in positively charged amino acids and five disulfide bonds within the cysteine-rich Agouti domain at the C-terminus (Fig1A). The cysteine-rich Agouti domain is the interaction site for MC1R and is common to both ASIP and its paralog AgRP (10). When resolved under reducing sample buffer conditions, the full-length glycosylated ASIP migrates similar in size to pro-ANP-FlagHis. Incubation with Corin generated faster migrating ASIP product(s) approximately half the size of the full-length ASIP (Fig1B). Consistent with pro-ANP cleavage, the intensity of the lower ASIP band also increased in a temperature dependent manner.

Next, we repeated the incubation of ASIP with twice as much soluble Corin and analyzed the products at 3 and 6 hours by two different methods. Corin digestion produced ASIP fragments in a time dependent manner resolved by SDS-PAGE under reducing sample conditions (Fig1C, green arrowhead). The intact masses of ASIP and the Corin digestion products were also measured by LC-MS using a method developed for quantitation of the NPPA prodomain (21). The mature full-length ASIP protein has an expected molecular weight of 12,044 Daltons (Da) with an unknown mass from N-glycosylation. The experimentally determined LC-MS intact mass for full-length glycosylated ASIP was found to be 14,177 Da, therefore the calculated glycosylation mass is 2,143 Da (Fig S1A). After 3 hours of incubation with Corin, a new ASIP product was generated with an intact mass of 6,170 (FigS1A, green arrow). This peak increased at the 6 hours timepoint and matched this glycosylated His23-Lys58 N-terminal region of ASIP. A second peak corresponding to the C-terminal region was not detected by this method. The deduced Lys58-Ser59 cleavage site is located at the start of the ASIP basic domain (Fig1A) and is highly conserved in mammals (FigS2). However the ASIP basic domain and the corresponding Corin cleavage site was absent from the paralog AgRP (8).

### Corin cleaves ASIP at the beginning of the basic domain

To confirm the location of the Corin cleavage site in the ASIP basic domain, we performed additional experiments to map the cleavage site. The Corin processed ASIP products have expected masses of 6.2 kDa for the N-terminus and 8 kDa for the C-terminus (Fig2A), however under reducing conditions the two products are poorly resolved. In a repeat experiment, Corin digested products were split into sample buffers with reducing and non-reducing conditions to increase the resolution of the ASIP cleaved fragments. Additionally, the C-terminal region contains more positively charged basic residues which will result in more intense Coomassie blue staining. Under non-reducing conditions, we observed a slower migrating ASIP product which stained more intensely with Coomassie blue, consistent with the deduced C-terminal cleavage product (Fig2B). In contrast, incubation of ASIP with soluble catalytic mutant form of Corin (Corin mut) did not result in ASIP cleavage. Using an antibody specific to the ASIP basic domain, we confirmed the identity of this C-terminal cleavage product migrating under reducing and non-reducing conditions (Fig2C).

To investigate the Corin cleavage site with Edman degradation or N-terminal protein sequencing, Corin and ASIP were incubated overnight resulting in near complete processing of ASIP (Fig2D). The N-terminal sequencing results were compared for ASIP alone, ASIP with Corin WT and ASIP with Corin mut (Fig2E and FigS1B-D). N-terminal sequencing of ASIP confirmed the signal peptide processed mature His23 sequence (FigS1B). Incubation with Corin WT produced two peaks for N-terminal sequencing, one consistent with the N-terminal region and the other beginning at Ser59 (Fig2E and FigS1C). Incubation with the Corin catalytic mutant resulted in a sequencing read of only the N-terminal region (FigS1D). Therefore the N-terminal sequencing results verify the Lys58-Ser59 cleavage site in human ASIP.

### ASIP peptides define the minimal substrate recognition site for Corin

To explore the requirements of the Corin cleavage site region, a series of ASIP peptides of different lengths spanning the Lys58-Ser59 cleavage were generated to establish a fluorogenic peptide assay (Fig3A). Each ASIP peptide contained an N-terminal AF488 reporter and a C-terminal QXL520 quencher. Intact peptides produce no signal but an endoproteinase cleavage event will release the C-terminal quencher resulting in fluorescent signal. Each peptide was incubated with soluble Corin, and the relative fluorescence was measured over time. Although the shortest 8-mer (LNKKSKQI) was not processed by purified Corin, incubation with purified Corin resulted in time dependent fluorescence for the 11-mer (ALNKK|SKQIGR) and 15-mer (IVALNKK|SKQIGRKA) ASIP peptides demonstrating that these peptides were sufficient for Corin substrate recognition and cleavage (Fig3B).

## Discussion

In this study we show that ASIP is a substrate for the Corin protease and identify the ASIP cleavage site at Lys58-Ser59 in the conserved basic domain. A single cleavage site was mapped using recombinant proteins analyzed by SDS-PAGE Coomassie blue stained gels and via western blot utilizing an epitope flanking the Lys58-Ser59 site. Edman degradation N-terminal sequencing confirmed the location of the Corin cleavage site. The Lys58-Ser59 site was further validated using ASIP fluorogenic peptide substrates and a purified soluble Corin to define a minimal substrate recognition region (ALNKK|SKQIGR). This region of ASIP is highly conserved in mammals consistent with an evolutionarily conserved Corin-ASIP cleavage event influencing coat color pigmentation in multiple species.

The MC1R-cAMP signaling pathway is one of the key nodes of regulation for pigmentation. Activation of this pathway in melanocytes produces the dark eumelanin pigment (Fig4A) and antagonism of this pathway via ASIP results in the lighter pheomelanin pigment (Fig4B). The ASIP co-receptor ATRN directly binds to the ASIP N-terminus and increases the effectiveness of ASIP (9). Therefore, Corin mediated cleavage at Lys58-Ser59 likely produces a C-terminal ASIP protein able to bind to MC1R via the Agouti cysteine-rich domain but without the ability to synergize with the ATRN receptor (Fig4C). As a result, co-expression of Corin and ASIP allows for intermediate antagonism to fine tune the degree of pheomelanin pigmentation.

**Figure 4.**
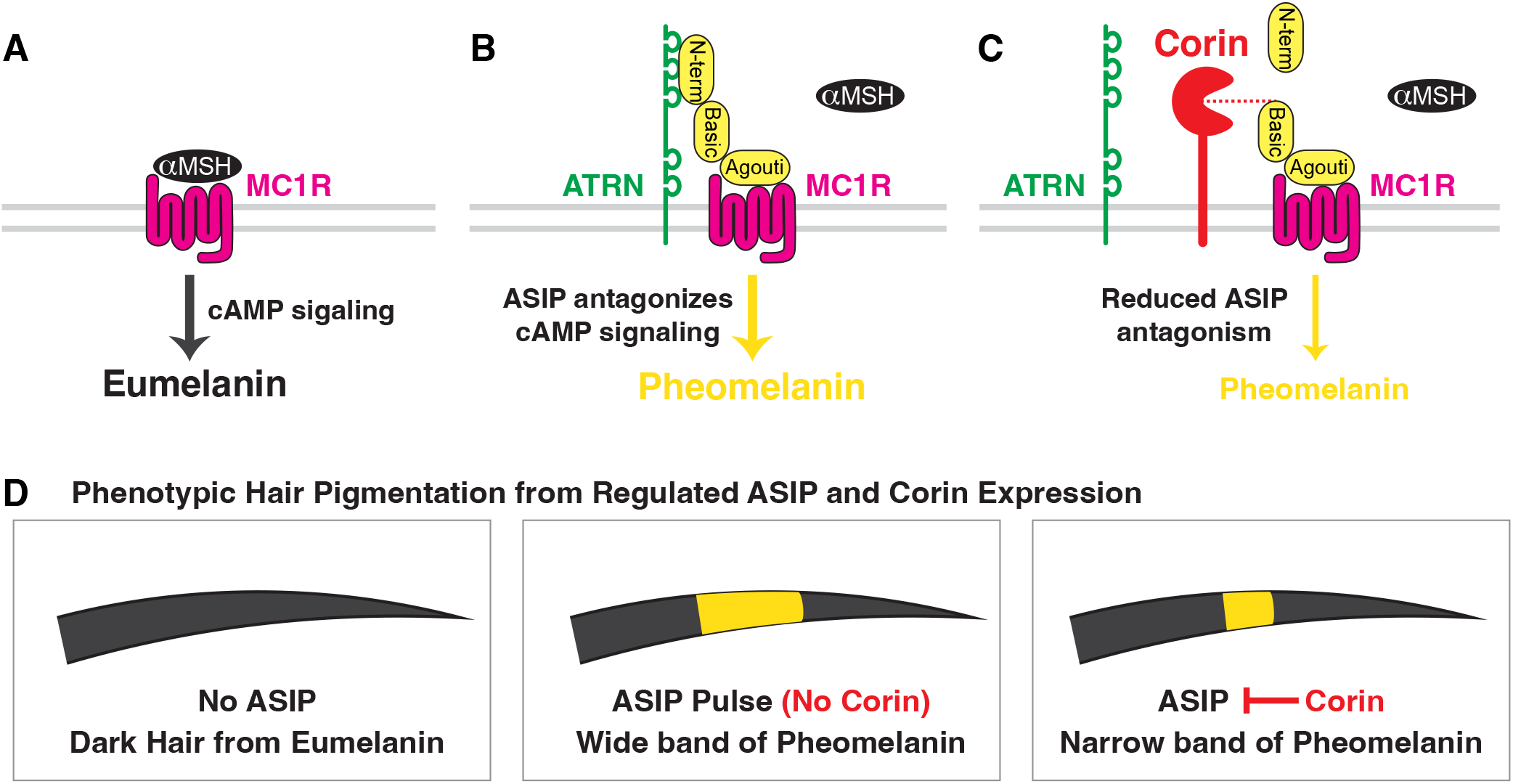
Model for Corin mediated inhibition of ASIP. A, α-MSH binds to MC1R resulting in cAMP signaling and eumelanin (dark pigment). B, ASIP binds to MC1R and co-receptor ATRN to inhibit α-MSH, resulting in pheomelanin (yellow pigment). C, Corin mediated ASIP cleavage removes the ATRN interaction domain, decreasing the effectiveness of ASIP to antagonize α-MSH/MC1R signaling. D, Phenotypic results for the hair shaft with variable ASIP and Corin expression. Corin cleavage of ASIP allows for fine tuning of the yellow pigmented region.

Transient expression of ASIP results in a subapical yellow band from the pigment switching from eumelanin to pheomelanin and loss of Corin expands the band of lighter pigmentation (Fig4D). Targeted catalytic mutation of Corin in the mouse results in a *wideband* phenotype and shows that enzymatic function is required for pigment switching (18). Several *wideband* Corin cat mutants have been described in various Corin extracellular domains with inferred loss of function. For instance, the *wideband* Golden cat Corin R795C mutant replaces a conserved arginine with an additional cysteine in the Scavenger receptor domain (15). It is worth noting that the corresponding human Corin variant R792C was also shown to have impaired proANP processing (21). Given the elaborate disulfide bond structure of Corin, many variants that affect disulfide bond formation or protein folding will likely impact both proANP and ASIP functions. However, it remains possible that specific domains of Corin are key for substrate recognition for either proANP or ASIP. This may be the case for the *wideband* Golden tiger Corin variant H587Y in the LDLR6 repeat (14). Preliminary in vitro characterization of this variant shows some evidence for diminished ASIP processing; however, it is unclear if this is due to reduced ASIP interaction or simply due to impaired Corin trafficking.

Natural variation at the Corin and ASIP loci has been shown to segregate with lighter coat color, including the seasonal white pelage of the white-tailed jackrabbit (22). In the wild *Peromyscus* deer mice, alternative promoters were shown to control Agouti/ASIP expression selectively in the dorsal and ventral regions and these regulatory regions were subject to evolutionary pressures from predation (23). At sites where the soil was lighter in color, lighter colored mice thrived with higher ASIP expression. Conversely, the classic loss of function ASIP mutations, or non-agouti represented by lowercase *a*, have reduced or absent bands of pheomelanin (7). For instance, the loss of function *a* allele for the black coated C57BL/6 mouse line results from a regulatory mutation in the first intron that blocks expression (24). In laboratory mice, a variety of missense and nonsense variants have been reported at the agouti locus, however no coat color missense variants have been observed at the Corin cleavage site (25). A hypermorphic ASIP variant that ablates the Corin cleavage but maintains function to regulate MC1R signaling would be expected to phenocopy the *wideband* Corin loss of function phenotype. Although an Agouti locus *wideband* mutant has not been reported in laboratory mice, a ΔSer48 coding variant near the Corin cleavage site has been described in *Peromyscus* deer mice from the Nebraska Sand Hills (26,27). Transgenic overexpression of the ΔSer48 allele resulted in a lighter coat color pigmentation, consistent with an ASIP hypermorph (28). It remains a possibility that removal of serine residue upstream from the conserved Lys58-Ser59 Corin cleavage site will impact substrate recognition or cleavage activity for the Corin enzyme.

## Acknowledgements

This work was supported by a research grant from Bayer AG within the Cardiovascular Disease Initiative at the Broad Institute. Dr. Ellinor was supported by grants from NIH (R01HL092577, R01HL157635, and R01HL139731), American Heart Association Strategically Focused Research Networks (18SFRN34110082), and the European Union (MAESTRIA 965286). We thank Michael Berne at Tufts University Core Facility for assistance with protein sequencing.

## Supplementary Figure List

Supplementary Figure 1. Identification of the ASIP cleavage site by LC-MS and N-terminal sequencing.

Supplementary Figure 2. Conservation of ASIP/Agouti at the Corin cleavage site.

