## Supplementary material for "Identification of a new Corin atrial natriuretic peptide-converting enzyme substrate: Agouti-signaling protein (ASIP)": FigS1

### Figure S1

**A** Uncleaved and Corin cleaved ASIP examined by LC/MS

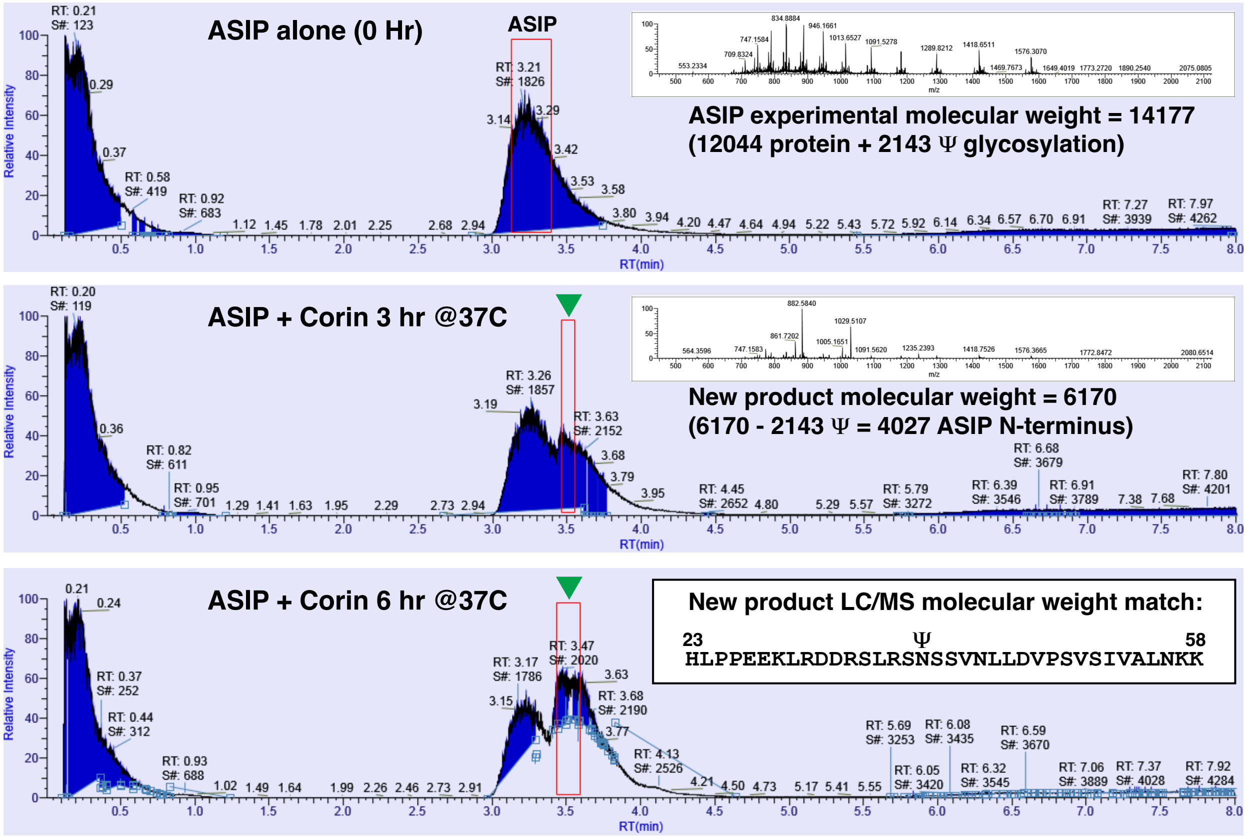


**B** ASIP N-terminal sequencing: HLPPEE

| Cycle | ASP | ASN | SER | GLN | THR | GLY | GLU | HIS | ALA | ARG | TYR | PRO | MET | VAL | TRP | PHE | ILE | LYS | LEU |
|---|---|---|---|---|---|---|---|---|---|---|---|---|---|---|---|---|---|---|---|
| 1 | 1.37 | 0.38 | 3.08 | 0.00 | 0.90 | 1.91 | 0.63 | **28.01** | 0.00 | 0.37 | 0.69 | 0.49 | 0.15 | 0.58 | 0.09 | 0.00 | 0.42 | 0.77 | 1.55 |
| 2 | 1.25 | 0.35 | 1.16 | 0.46 | 0.60 | 1.86 | 0.54 | 5.36 | 1.42 | 1.15 | 0.59 | 1.04 | 0.20 | 1.01 | 0.00 | 0.00 | 0.53 | 0.76 | **54.74** |
| 3 | 3.03 | 0.94 | 1.84 | 0.70 | 0.50 | 2.20 | 0.74 | 1.59 | 1.55 | 2.33 | 0.72 | **47.66** | 0.51 | 1.46 | 0.00 | 0.46 | 0.66 | 1.57 | 14.24 |
| 4 | 2.96 | 1.01 | 2.28 | 0.84 | 0.53 | 1.73 | 0.94 | 0.62 | 1.28 | 2.02 | 0.62 | **35.03** | 0.52 | 1.34 | 0.08 | 0.32 | 0.83 | 2.04 | 3.63 |
| 5 | 1.60 | 1.26 | 2.27 | 0.83 | 0.35 | 1.88 | **25.43** | 0.17 | 1.74 | 1.64 | 0.56 | 18.14 | 0.62 | 2.04 | 0.09 | 0.26 | 0.87 | 2.75 | 2.87 |
| 6 | 1.14 | 1.53 | 2.57 | 1.03 | 0.50 | 2.18 | **39.48** | 0.23 | 2.29 | 2.00 | 0.66 | 8.37 | 0.56 | 2.51 | 0.00 | 0.40 | 1.04 | 3.10 | 2.92 |

**C** ASIP +Corin WT N-terminal sequencing: HLPPEE and SKQIGR

| Cycle | ASP | ASN | SER | GLN | THR | GLY | GLU | HIS | ALA | ARG | TYR | PRO | MET | VAL | TRP | PHE | ILE | LYS | LEU |
|---|---|---|---|---|---|---|---|---|---|---|---|---|---|---|---|---|---|---|---|
| 1 | 1.69 | 2.32 | **27.44** | 0.00 | 1.64 | 1.62 | 0.50 | **25.68** | 3.51 | 1.62 | 1.26 | 0.20 | 0.34 | 0.74 | 0.14 | 0.00 | 0.81 | 18.09 | 0.00 |
| 2 | 1.46 | 0.53 | 9.82 | 0.13 | 0.51 | 1.45 | 0.49 | 6.91 | 5.57 | 1.77 | 0.81 | 2.78 | 0.41 | 2.07 | 0.09 | 0.00 | 0.59 | **36.32** | **67.83** |
| 3 | 3.35 | 1.44 | 3.28 | **22.70** | 0.00 | 1.56 | 4.72 | 1.55 | 2.26 | 2.66 | 0.56 | **39.98** | 0.40 | 1.74 | 0.08 | 0.16 | 0.48 | 16.99 | 12.61 |
| 4 | 3.49 | 1.58 | 5.33 | 17.08 | 0.00 | 1.93 | 4.40 | 0.98 | 1.69 | 3.09 | 0.72 | **46.74** | 0.58 | 1.76 | 0.00 | 0.30 | **24.48** | 10.91 | 5.47 |
| 5 | 1.69 | 1.31 | 3.31 | 6.06 | 0.45 | **13.42** | **25.71** | 0.65 | 2.44 | 2.62 | 0.67 | 18.99 | 0.53 | 2.28 | 0.08 | 0.16 | 15.18 | 7.56 | 3.22 |
| 6 | 1.42 | 1.45 | 3.22 | 3.49 | 0.79 | 12.58 | **41.44** | 0.38 | 2.99 | **11.63** | 0.71 | 9.72 | 0.53 | 2.57 | 0.11 | 0.24 | 7.42 | 6.14 | 3.07 |

**D** ASIP +Corin mut N-terminal sequencing: HLPPEE

| Cycle | ASP | ASN | SER | GLN | THR | GLY | GLU | HIS | ALA | ARG | TYR | PRO | MET | VAL | TRP | PHE | ILE | LYS | LEU |
|---|---|---|---|---|---|---|---|---|---|---|---|---|---|---|---|---|---|---|---|
| 1 | 2.65 | 1.05 | 13.63 | 0.00 | 2.26 | 6.26 | 1.44 | **24.31** | 1.94 | 2.57 | 1.93 | 0.86 | 0.44 | 1.46 | 0.16 | 1.07 | 1.73 | 1.36 | 4.30 |
| 2 | 1.77 | 0.96 | 3.11 | 1.11 | 1.28 | 6.39 | 1.37 | 5.14 | 3.44 | 3.41 | 1.76 | 2.35 | 0.64 | 2.32 | 0.00 | 1.40 | 1.42 | 1.25 | **55.50** |
| 3 | 3.72 | 2.03 | 3.02 | 1.36 | 1.23 | 6.62 | 1.66 | 1.52 | 2.28 | 8.25 | 1.54 | **32.12** | 0.63 | 2.14 | 0.00 | 1.18 | 1.21 | 1.82 | 12.59 |
| 4 | 3.59 | 1.86 | 6.62 | 1.47 | 0.90 | 6.02 | 1.97 | 0.72 | 2.04 | 4.83 | 1.49 | **28.31** | 0.64 | 2.29 | 0.13 | 1.05 | 1.50 | 2.53 | 5.28 |
| 5 | 3.19 | 2.22 | 6.09 | 1.93 | 1.23 | 6.96 | **24.80** | 0.82 | 3.46 | 4.81 | 1.68 | 18.16 | 0.75 | 3.94 | 0.00 | 1.07 | 1.76 | 3.39 | 5.22 |
| 6 | 2.64 | 2.10 | 5.81 | 1.75 | 1.33 | 5.77 | **33.91** | 0.67 | 3.64 | 3.85 | 1.44 | 7.51 | 0.66 | 3.61 | 0.06 | 1.04 | 1.71 | 3.39 | 4.34 |

**Supplementary Figure 1. Identification of the ASIP cleavage site by LC-MS and N-terminal sequencing.** A, Time dependent ASIP cleavage by Corin resolved by LC-MS. Molecular weights for the intact purified ASIP and a new Corin dependent product (green arrowhead) are consistent with ASIP cleavage at Lys58-Ser59. B-D, N-terminal sequencing results for ASIP, ASIP +Corin WT and ASIP +Corin mut. Edman degradation products for each cycle are resolved by HPLC-MS and peaks for each amino acid were quantitated. Results are plotted in a tabular view of the peak intensity values for six cycles. Relative intensities are shaded in green. Multiple peaks in the ASIP +Corin WT condition demonstrate both mature ASIP (beginning at His23) and the ASIP cleavage product (beginning at Ser59, shown in red).
