## Supplementary material for "Identification of a new Corin atrial natriuretic peptide-converting enzyme substrate: Agouti-signaling protein (ASIP)": FigS2

### Figure S2

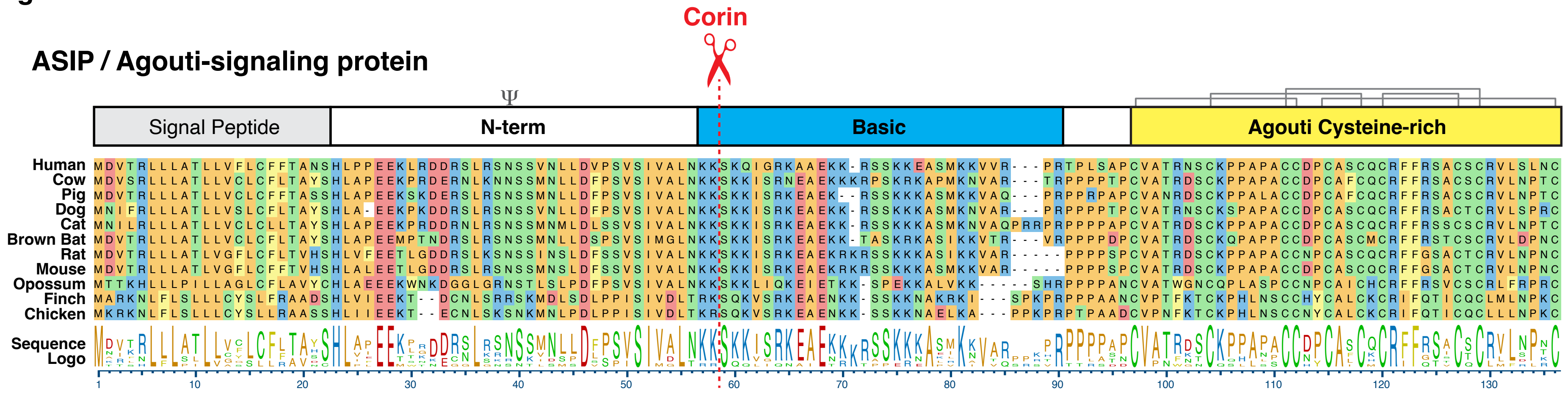


**Supplementary Figure 2. Conservation of ASIP/Agouti at the Corin cleavage site.** Clustal Omega alignment of ASIP proteins from Human (P42127), Cow (Q29414), Pig (Q6ZYM3), Dog (Q5UK76), Cat (Q865F0), Brown Bat (G1NUR0), Rat (Q99JA2), Mouse (Q03288), Opossum (F6YH41), Zebra Finch (A0A674GJ99) and Chicken (C9K601). Agouti Cysteine-rich domain disulfide bonds and the mammalian N-glycosylation site are shown above the protein domains. The Corin cleavage site at human ASIP Lys58-Ser59 is highly conserved.
